# Homogeneously crosslinked *in situ* hydrogel enclosing high-density human-cancer cells promotes vascularized *in vivo* tumor modeling for immune cell therapy

**DOI:** 10.1101/2024.12.17.628981

**Authors:** Ziqi Huang, Yip Ming Tsun, Chao Liang, Zhenzhen Wu, Theo Aurich, Lu Liu, Rio Ryohichi Sugimura, Sang Jin Lee

## Abstract

Cancer models in animal studies play a central role in cancer research, particularly in investigating vascularized tumor tissues for the validation of immune cell therapies. However, xenografts relying solely on cancer cells are ineffective for optimal tumor tissue formation. Additionally, tumor modeling using hydrogels with cancer cells to promote vascularization often leaves behind residual biomaterials that inhibit integration with surrounding tissues. To address these issues, we utilized a straightforward *in vivo* vascularized tumor modeling method with a completely degradable, crosslinker-free carboxymethyl chitosan (CMCTS)/oxidized hyaluronic acid (OHA) hydrogel that encapsulates high-density human cancer cells for *in situ* injection. The CMCTS/oHA hydrogel was fully degraded within 3 weeks, enabling three-dimensional (3D) cell condensation *in vitro*. 2 weeks after subcutaneous injection in mice, solid tumors formed, with native host vasculature infiltrating the transplanted human cancer cells, confirming spontaneous hydrogel degradation. Following this, human macrophages were administered via tail vein injection, enhancing the accumulation of mouse immune cells in the humanized tumor twofold and showing murine macrophages adjacent to the vasculature. This study thus provides proof-of-concept for a facile and fully vascularized humanized tumor model in mice for validating immune cell therapies.

**HIGHLIGHTS:**

- The oHA was prepared using sodium periodate treatment, which facilitated the formation of *in situ* CMCTS/oHA hydrogels
- CMCTS/oHA hydrogels completely degraded within a short period, allowing for 3D cell condensation
- High-density cell-laden CMCTS/oHA hydrogels were injected subcutaneously in mice, resulting in the generation of a vascularized solid tumor
- The transplanted therapeutic cell was observed to adhere to the tumor tissue through the bloodstream

## 1 INTRODUCTION

Cancer is one of the most formidable challenges in global health care, accounting for nearly one in six deaths worldwide [1]. A significant proportion of cancer research efforts— encompassing basic science, translational medicine, and preclinical trials—frequently fail to yield effective treatment outcomes, resulting in a tremendous waste of time, financial resources, and labor-intensive efforts [2]. Although many scientists and medical doctors have made important contributions to cancer research, progress has been significantly delayed due to the complexities of cancer modeling, which hampers the testing of newly investigated drugs and therapeutic tools both *in vitro* and *in vivo* [3, 4].

To conduct more rigorous cancer research, establishing effective tumor models is essential for advancing the field and minimizing unnecessary resource wastage. So far, traditional xenograft models have been widely used, involving the transplantation of cancer cells into animal hosts. However, these models are often unsuitable for studying early-stage cancer due to their low engraftment rates and limited metastatic potential [5, 6]. Indeed, the limitations of traditional xenograft models result in poor predictive power when evaluating anti-cancer therapies for future clinical use [7]. Indeed, vascularized tumor models are particularly critical for studying tumor progression, as tumors rely on angiogenesis for growth and metastasis. Furthermore, immunotherapy and drug delivery require functional vascularized models to assess efficacy through intravenous injection [8]. Generally, engineered *in vitro* tissues are vascularized by co-culturing endothelial cells (ECs) and human tumor cells; however, these models often lack specificity and fail to form a perfusable network, limiting their relevance [9]. While *in vitro* vascularization strategies utilizing hydrogel pre-patterning techniques have demonstrated the ability to generate perfusable single vessels [10], they struggle to produce capillary-sized channels due to difficulties in endothelialization, making it challenging to mimic native complex vascular beds [11]. To address this issue, vascularized tumor channels have been developed using lab-on-a-chip technology, but this approach often involves time-consuming processes and frequent failures *in vitro* to create viable channels [12, 13]. Regrettably, vascularized tumor models in animals remain indispensable, as the mature primary vascular network of tumors provides comprehensive and reliable feedback in the development of effective cancer therapies.

In current *in vivo* xenograft models, cancer cells are inoculated into animal subjects; however, these models often suffer from unexpected cell loss and dispersion post-transplantation, as the transplanted cells cannot sustain themselves without a supportive matrix [14].

To address these issues, various hydrogel systems have been employed to enhance *in vivo* tumor modeling, including ultraviolet (UV) crosslinking [15], thermal crosslinking [16] and other chemical crosslinking systems [17]. While these systems aim to provide a supportive environment for tumor growth and vascularization, they face significant challenges. For instance, UV-based and thermo-responsive hydrogels can negatively impact cell viability and functionality due to the prolonged conditions required for gelation [18]. Furthermore, these crosslinking methods may result in poor hydrogel degradability, limiting tissue integration and vascularization with native host tissue [19]. The specialized equipment required for these processes, such as UV crosslinking machines, along with time-consuming protocols, can introduce variables that complicate reproducibility and scalability [20]. Thus, new solutions should be investigated for a clear vascularized tumor model *in vivo*. Therefore, new solutions must be explored to establish a reliable vascularized tumor model *in vivo*. Unfortunately, a facile hydrogel capable of delivering cancer cells while simultaneously promoting vascularization through spontaneous degradation has yet to be clearly established in cancer research.

Herein, we engineered a polysaccharide-based *in situ* hydrogel for encapsulating high-density human cancer cells through simple injection to model an *in vivo* vascularized tumor tissue. We utilized carboxymethyl chitosan (CMCTS) and oxidized hyaluronic acid (oHA) for the local delivery of cancer cells as a cell-holding approach before injection. In our previous studies, Lee *et al.* introduced water-soluble glycol chitosan (gC) and oHA-based self-healing hydrogels, which were validated for *in situ* therapy in rat calvarial [21] and spinal cord injury model [22]. Building on these motifs, we anticipated the use of self-assembling hydrogels that are volume-controllable and easy to fabricate without requiring crosslinking agents. This approach offers a user-friendly and rapidly degradable alternative to facilitate the formation of vascularized tumor models within the animal body. Additionally, we envisioned that the hydrogel would provide structural support and adhesion sites for tumor cells post-injection, while its rapid degradation promotes integration with the native tissue microenvironment and vascular infiltration into the newly formed tumor tissue. A detailed schematic illustration of this approach is depicted in Fig. 1.

**Fig. 1.**
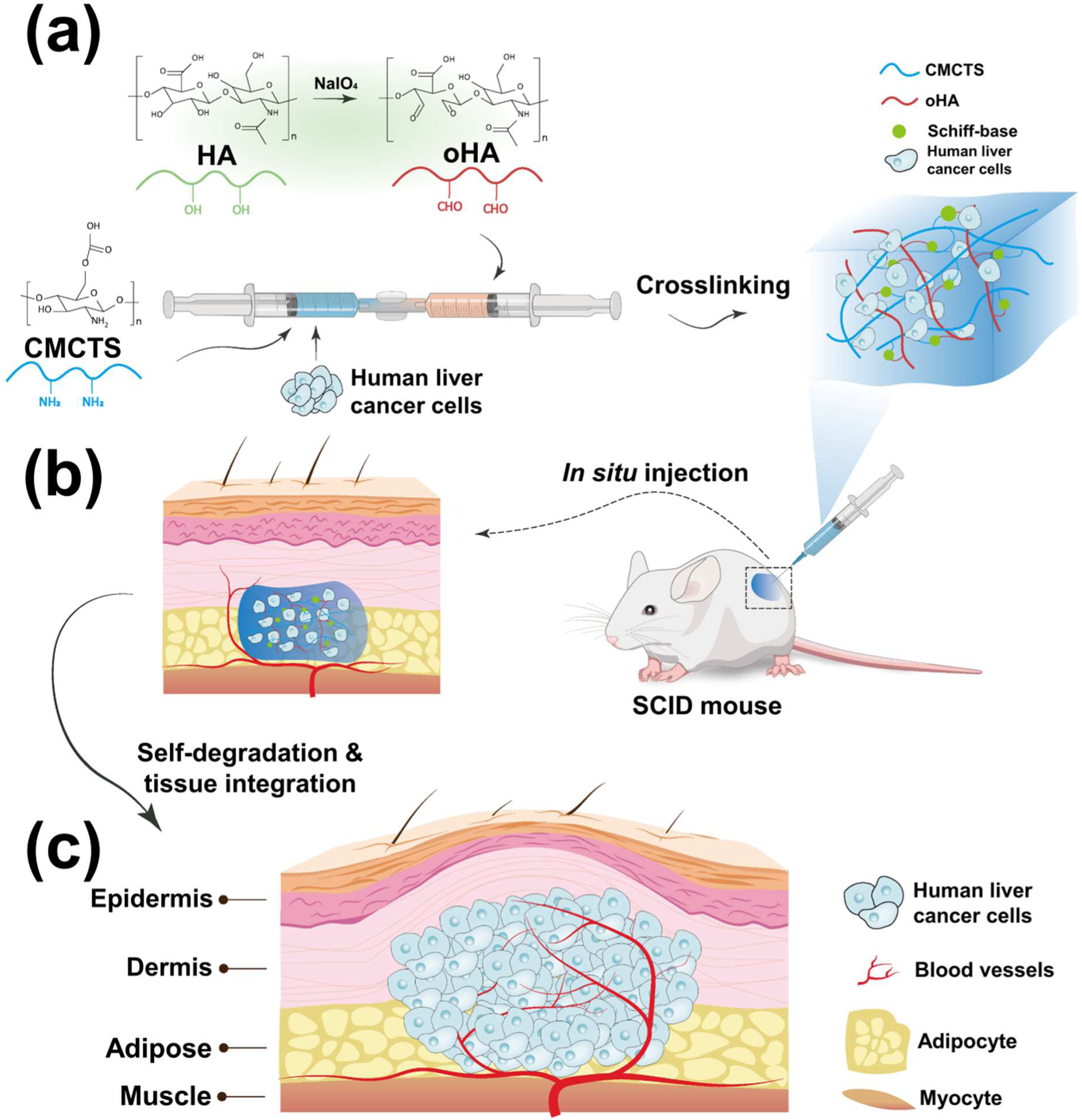
Schematic illustration of the preparation procedures of high-density human liver cancer cells-encapsulated CMCTs/oHA injectable hydrogel and their *in vivo* application. CMCTS/oHA hydrogels were homogeneously mixed with human liver cancer cells to form an injectable *in situ* hydrogel (a). This hydrogel was immediately injected subcutaneously in mice (b), leading to the formation of a healthy vascularized tumor model through the integration of cells released from the hydrogel after degradation (c)

This work aims to investigate whether (1) the hydrogel can be rapidly degraded in mice after injection, (2) a tumor model can be spontaneously generated without further intervention, (3) a robust vasculature can be established, leading to the development of condensed solid tumors in the mice, and (4) the potential efficacy of immune cell treatment can be assessed. To achieve this, we analyzed the physicochemical properties of the hydrogel and conducted studies on *in vitro* three-dimensional (3D) cell cultures, as well as tissue evaluation and immune cell therapy potential in animal models

## 2 Experimental Section

### 2.1 Materials

Hyaluronic acid powder with high molecular weight (HA, 1000 kDa, Cosmetic & Food Grade) was purchased from The Pharmers Market (Manchester, United Kingdom). O-carboxymethylated chitosan (CMCTS, Cat. No. sc-358091), with a deacetylation degree of ≥ 90%, was obtained from Santa Cruz Biotechnology (Santa Cruz, CA, USA). Deuterium oxide (D₂O, Cat. No. 151882, isotopic purity of 99.9 atom % D) was supplied by Sigma-Aldrich (St. Louis, MO, USA). Luer-lock tip syringes (1.0 mL, Cat. No. 309628) were acquired from BD Biosciences (Becton Dickinson, USA), while tunnel mixers (Cat. No. MT-SEM01) for connecting two syringes were purchased from NOBAMEDI Co., Ltd. (Yongin, Republic of Korea).

### 2.2 Preparation of oHA and CMCTS/oHA hydrogels

First, oHA was prepared by sodium periodate oxidation, as described in previous literature [21, 22]. In brief, 360 mL of distilled water was used to dissolve 3.8 g of HA. The HA solution was then oxidized in the dark at room temperature by adding 40 mL of 5.0 mmol NaIO₄. After about six hours, excess ethylene glycol (1 mL) was added to the HA solution to terminate the reaction for 30 minutes. Subsequently, dialysis packs (Cat. No. JHC0095, 8000-14000 MWCO, BioMomei, China) were used to dialyze the solution for a week, with deionized water changed twice a day to eliminate any residual products. Finally, the oHA composite was obtained by lyophilization over seven days and stored at -20 °C for further use.

The CMCTS/oHA hydrogels were prepared as described in previous literature. Briefly, CMCTS was dissolved in phosphate-buffered saline (PBS) to form a 4% (w/v) macromer solution, and the synthesized oHA composite was similarly dissolved in PBS to form a 3% (w/v) solution. CMCTS/oHA hydrogels were then formed by thoroughly mixing 4% CMCTS and 3% oHA in an 80:20 ratio at room temperature for 3 minutes using syringes with a tunnel mixing tool.

### 2.3 Characterization of CMCTS/oHA hydrogels

#### 2.3.1 ^1^H-Nuclear magnetic resonance (^1^H-NMR) spectra and fourier-transform infrared spectroscopy (FT-IR)

To confirm the successful synthesis of aldehyde groups, the produced oHA was dissolved in D2O and then determined using a ^1^H-NMR spectrometer (Bruker Avance 400, Bruker Corporation, Billerica, MA, USA). The CMCTS compounds and oHA composites, and lyophilized CMCTS/oHA hydrogels were characterized by FT-IR spectrometer (Spectrum Two™, PerkinElmer, Norwalk, CT, USA) within a wavenumber range of 400 to 4000 cm⁻¹.

#### 2.3.2 *In vitro* degradation test

CMCTS/oHA hydrogels were prepared as described above. After lyophilization for 72 hours, the initial weight (W₀) of the CMCTS/oHA composite was measured. The degradation rate of the CMCTS/oHA composites was assessed by consistent shaking in PBS using a shaking incubator (Stuart Orbital Incubator S150, Staffordshire, UK) at 37°C with a speed of 80 rpm. Fresh PBS was carefully replaced every 24 hours. Samples were collected at predetermined time points (1, 2, 3, 5, 7, and 14 days; n = 4 for each time point) and lyophilized for 72 hours before being photographed and weighed (Wₜ). The degradation rate was defined using the following formula.

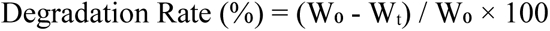

#### 2.3.3 Rheology test

The rheological properties of CMCTS/oHA hydrogels were examined using a rheometer (HAAKE MARS 40, Thermo Scientific, Waltham, MA, USA) at room temperature. As mentioned before, 4% CMCTS solution was mixed with 3% oHA solution by syringes with 2-side tunnel mixer to synthesize hydrogel for further measurements. Afterwards, the prepared hydrogels were put on the rheometer plate with geometrical parameters of 1 mm gap and 8 mm diameter plate-plate configuration to examine the viscoelastic behavior. All samples were evaluated for storage elastic modulus (G′) and loss modulus (G″) under controlled circumstances of room temperature and steady shear flow technique. A constant strain of 1% was applied while the frequency sweep was conducted in the 0.1–10 Hz range throughout the whole testing procedure.

### 2.4 Human liver cancer cells encapsulated CMCTS/oHA hydrogels

#### 2.4.1 Preparation of HepG2 cells and Huh-7 cells

HepG2 cells (Cat. No. HB-8065, ATCC, Manassas, Virginia, USA) were cultured in Eagle Minimum Essential Medium (EMEM, Cat. No. 130-2003, ATCC, Manassas, Virginia, USA) containing 10% Fetal Bovine Serum (FBS, Cat. No. 10270106, Life Technologies, Carlsbad, California, USA) and 1× Penicillin–Streptomycin (Cat. No. 15140-122, Gibco, Waltham, Massachusetts, USA) in a 150 mm cell culture dish (Cat. No. CS016-0129, ExCell Bio Inc., Shanghai, China).

Huh-7 cells (Cat. No. CVCL_0336, Cellosaurus, Geneva, Switzerland) were cultured in Dulbecco’s Modified Eagle Medium (DMEM, Cat. No. 11995065, Life Technologies, Carlsbad, California, USA) containing 10% FBS (Cat. No. 10270106, Life Technologies, Carlsbad, California, USA) and 1× Penicillin–Streptomycin (Cat. No. 15140-122, Gibco, Waltham, Massachusetts, USA) in a 150 mm cell culture dish (Cat. No. CS016-0129, ExCell Bio Inc., Shanghai, China).

The medium was replaced twice a week to maintain optimal growth conditions. Cell pellets were obtained when the cultures reached approximately 80-90% confluence. After aspirating the medium and washing the cells once with PBS (Cat. No. 20012027, Life Technologies, Carlsbad, California, USA), Trypsin-EDTA (0.25%) (Cat. No. 25200072, Gibco, Waltham, Massachusetts, USA) was added to the culture flasks and incubated for 8 minutes at 37 °C until the cells were fully detached from the plates. The trypsin and cell suspension were neutralized by adding an equal volume of complete medium, and then centrifuged at 1800 rpm for 5 minutes. The cells were then used for the CMCTS/oHA hydrogel combination, as described in section 2.4.2.

#### 2.4.2 Encapsulation of HepG2 or Huh-7 cells in CMCTS/oHA hydrogels

To encapsulate the cells within CMCTS/oHA hydrogels, HepG2 cell pallets and Huh-7 cell pallets were mixed thoroughly with 4% CMCTS macromer solution at a density of 150 x 10^6 cells/ml, and then CMCTS/cells suspension was directly mixed with oHA. The detailed hydrogel preparation procedure was the same as above in Section 2.2. After vigorous mixing, 50 μL of HepG2 or Huh-7 encapsulated CMCTS/oHA hydrogels were loaded on a 48-well plate. Each well contains 700 μL of high sugar medium and incubated at 37 °C in a humidified 5% CO2 incubator for 3 days, with the medium being changed once a day and imaging to observe their morphologies. The macroscopic morphology of cell-laden hydrogels was determined by the camera and stereoscope (SMZ18 Stereo Zoom Microscope, Nikon Instruments Inc., USA). Additionally, the microstructure of hydrogels was investigated using a Scanning Electron Microscope (SEM, Cat. No. SU-1510, Hitachi High-Technologies, Tokyo, Japan) after the 72-h lyophilization and sputter-coating with Au.

#### 2.4.3 Characterization of HepG2 or Huh-7 aggregates from CMCTS/oHA hydrogels

Hydrogels were collected on the third day and a Hoechst 33342/Propidium Iodide (PI) Double Staining kit (Cat. No. BL116A, Biosharp, Hefei, China) was used to test the viability of HepG2 and Huh-7 cells. Remaining samples were embedded and frozen at -80 °C overnight, and then cryosectioned using a cryotome (Cat. No. Leica CM 1860 UV, Microsystems Inc., Wetzlar, Germany). Sections were then treated with F-actin fluorescence staining. Samples were washed with PBS, fixed in 3.7% formaldehyde for 20 minutes, permeabilized with 0.5% Tween 20 for 10 minutes, blocked with 1% BSA for 30 minutes, stained with Actin (1:100) for 30 minutes, and treated with DAPI for 3 minutes, all at room temperature with PBS washes between steps. Slides were visualized through a confocal laser scanning microscope (LSM 900, ZEISS, Germany).

### 2.5 *In vivo* study

#### 2.5.1 Animal procedures

All animal studies were conducted under the animal research protocol (No. 22-268) approved by the Committee of the Use of Live Animals in Teaching and Research (CULATR) at the University of Hong Kong. The studies adhered to the Animals (Control of Experiments) Ordinance (Hong Kong) and guidelines from the Centre for Comparative Medical Research (CCMR), Li Ka Shing Faculty of Medicine, The University of Hong Kong.

Male BALB/c (CCMR), Male severe combined immunodeficiency, C.B-17/Icr-scid (SCID) mice, and Female NOD.Cg-Prkdc^scid^Il2rg^tm1Wjl^/SzJ (NSG) (21-25g, 6-8 weeks old) were used for the animal experiments. The mice were obtained from the Experimental Animal Center of the University, with access to food and water ad libitum, and were maintained under pathogen-free conditions. All procedures were performed under anesthesia using intraperitoneal injections of ketamine/xylazine mixtures (100 mg/kg ketamine (Cat. No. 013004, AlfaMedic Ltd.) + 10 mg/kg xylazine (Cat. No. 013006, AlfaMedic Ltd.). Anesthesia was confirmed by checking body reflexes, and eye ointment (Duratears® ointment, Cat. No. 05686, Alcon, Fort Worth, TX, USA) was applied to prevent blindness. The subcutaneous injection sites on the mice were decontaminated with alternating applications of 70% alcohol and povidone-iodine (Betadine®) using sterile cotton swabs. After injection, the mice were placed in an intensive care unit (ICU) until they fully recovered, after which they were returned to their original housing location. Seven days post-injection, the mice were euthanized by an overdose of Pentobarbital (250 mg/kg) (Dorminal®, Cat. No. 013003, AlfaMedic Ltd.) administered intraperitoneally.

#### 2.5.2 Injection of HepG2 or Huh-7 cells encapsulated CMCTS/oHA hydrogels in mice subcutaneously for vascularized tumor formulation

HepG2 or Huh-7 cells encapsulated hydrogels with the same cell concentration as in Section 2.4 were prepared. Cell-laden CMCTS/oHA hydrogels were then injected into the left and right backs of the mice, with 500 μL volumes per site using a syringe with a 23-gauge needle. Mice were sacrificed at the indicated time intervals at 1, 2, and 4 weeks. The injected area was carefully sectioned and photographed.

#### 2.5.3 Histological analysis *in vitro* and *in vivo*

The harvested tissues were embedded in paraffin after dehydration through a series of ethanol, ethanol-xylene, and xylene solutions. After embedding, the paraffin block was sectioned at 15 μm thickness using a rotary microtome (order number: RM2155, Leica Microsystems, Wetzlar, Germany). Sections were then deparaffinized, rehydrated, and stained with hematoxylin (Cat. No. SH4777, Harris Hematoxylin, Cancer Diagnostic Inc., Durham, NC, USA) and eosin (Cat. No. CS701, Dako, Glostrup, Denmark) (H&E), alcian blue (pH 2.5) with 0.1% nuclear fast red (Cat. No. PH1082, Scientific Phygene®, Fuzhou, China), and 0.5% safranin O (Cat. No. 02782-25, Polysciences Inc., Warrington, PA, USA) with fast green (Cat. No. PH1852, Scientific Phygene®), Wright-Giemsa (Cat. No. PH1793, Scientific Phygene®). The stained slides were observed using an optical microscope (Cat. No. Eclipse LV100POL, Nikon, Tokyo, Japan) equipped with a CCD camera (Cat. No. DS-Ri1, Nikon, Tokyo, Japan).

#### 2.5.4 IVIS Imaging

Bioluminescent images of tumors were captured using an *in vivo* bioluminescent imaging system (IVIS, Cat. No. Lumina X5, PerkinElmer Inc., Waltham, MA, USA). Mice received intraperitoneal injections of D-luciferin (Cat. No. LUCK-100, Gold Biotechnology, Inc., Olivette, MO, USA) at a dose of 150 mg/kg body weight under anesthesia, and placed in a prone position in XIC-3 animal isolation chamber (Cat. No. 123997, PerkinElmer). Images were collected with the XIC-3 chamber connected to the IVIS Spectrum and the photons emitted by the tumor and its surroundings were quantified using Living Image® software (PerkinElmer Inc., Waltham, MA, USA). Finally, all collected images were normalized and the average luminescence intensity of each mouse abdomen was calculated and reported.

#### 2.5.5 Immunofluorescence (IF) against anti-human CD45, anti-human GPC3, anti-mouse CD31, and anti-mouse F4/80 antibodies

For IF staining, antigen retrieval was performed through 1x antigen retrieval buffer (pH 6) (Cat. No. ab93684, Abcam., Cambridge, Cambridgeshire, United Kingdom). Deparaffinized slides in the chamber were then incubated in the microwave for 15 min prior to IF staining. Sections were then blocked using the blocking buffer: 3% Bovine Serum Albumin (BSA) (Cat. No. 2391492, Gibco, Waltham, Massachusetts, United States) in 0.1% Triton X-100 (Cat. No. T8787-100mL, Sigma Aldrich, St. Louis, Missouri, United States) in PBS for 15 minutes. After treatment, slides were incubated with primary antibodies without washing overnight at 4 °C. The other day, slides were washed with 1 x PBS + Triton X-100 (Cat. No. T8787-100mL, Sigma Aldrich, St. Louis, Missouri, United States) (PBST) for 3 times carefully, and incubated with secondary antibodies at room temperature for 2 hours, followed by DAPI staining (Cat. No. 564907, BD Biosciences, Franklin Lakes, New Jersey, United States) for 5 minutes. Slides were mounted with Anti-Fade fluorescent mounting medium (Cat. No. ab104135, Abcam, Cambridge, United Kingdom) and visualized through Nikon Confocal Microscope System AX / AX R (Chinetek Scientific (China) Limited, Hong Kong, China). The information of primary and secondary antibodies is below:

The primary antibodies: 1) Rabbit anti-human GPC3 (1:500) (Cat. No. ab207080, Abcam., Cambridge, Cambridgeshire, United Kingdom), 2) Rabbit anti-mouse CD31(1:2000) (Cat. No. ab8216, BD Biosciences, Cambridge, Cambridgeshire, United Kingdom), 3) Mouse anti-human CD45 from Abcam (Cat. No. ab10558, BD Biosciences, Cambridge, Cambridgeshire, United Kingdom), and 4) Rat anti-mouse F4/80 from ThermoScientific (Cat. No.14-4801-82, ThermoFisher Scientific, Waltham, Massachusetts, United States). Secondary antibodies: 1) Donkey anti-Mouse IgG (H+L) Highly Cross Adsorbed Secondary Antibody, Alexa Fluor 488 (Cat. No. A21202, ThermoFisher Scientific, Waltham, Massachusetts, United States), 2) Goat anti-Rabbit IgG (H+L) Cross-Adsorbed Secondary Antibody, Alexa Fluor 568 (Cat. No. A11036, ThermoFisher Scientific, Waltham, Massachusetts, United States), 3) Goat anti-Rat IgG (H+L) Cross-Adsorbed Secondary Antibody, and 4) Alexa Fluor 647 (Cat. No. A21247, ThermoFisher Scientific, Waltham, Massachusetts, United States).

### 2.6 Peripheral Blood Mononuclear Cell (PBMC) and CD14+ Monocyte Isolation, and Macrophage Differentiation

#### 2.6.1 PBMC Isolation from Whole Blood

Primary human monocytes were isolated and purified from PBMCs in the buffy coat obtained from healthy donors (Hong Kong Red Cross Blood Transfusion Service, Hong Kong) with the assistance of Professor C.S. Lau’s group, using Ficoll-Paque PLUS density gradient media (Cat No. 17144002, Cytiva, Marlborough, Massachusetts, United States) according to the manufacturer’s instructions. Briefly, 40 mL of blood was diluted 1:1 with PBS and carefully overlaid onto 15 mL of Ficoll, ensuring the layers did not mix. After centrifugation at 1000 x g for 30 minutes at room temperature with the brake turned off, plasma, PBMCs, Ficoll-Paque PLUS, and red blood cell layers were separated. The PBMC layer was then harvested and washed with 50 mL of PBS, followed by centrifugation at 300 x g for 5 minutes at room temperature with the brake on. Red blood cells were lysed by adding ACK Lysing Buffer (Cat No. A1049201, Gibco, Waltham, Massachusetts, United States) for 10 minutes at room temperature. The cells were then topped up with 50 mL of PBS and centrifuged at 300 x g for 5 minutes at room temperature. The supernatant, containing the lysed red blood cells and ACK lysis buffer, was removed. Finally, PBMC pellets were resuspended in 10 mL of PBS and counted using an automated cell counter.

#### 2.6.2 CD14+ monocytes isolation from PBMC

CD14+ monocytes were isolated from PBMC by CD14 MicroBeads, human (Cat No. 130-097-052, Miltenyibiotec, Bergisch Gladbach, North Rhine-Westphalia, Germany) according to manufacturer’s instructions. Briefly, cell pellets were resuspended in 80 µL of MACS buffer (0.5% BSA + 2 mM EDTA in PBS) per 10⁷ total cells. 20 µL of CD14 MicroBeads per 10⁷ total cells and incubate for 15 minutes at 2-8 °C. Cells were passed through the LS Column (Cat No. 130-122-729, Miltenyibiotec, Bergisch Gladbach, North Rhine-Westphalia, Germany) and labeled CD14+ monocytes were flushed out by the plunger.

#### 2.6.3 Macrophage Differentiation from CD14+ Monocytes

Isolated monocytes were resuspended in macrophage differentiation medium, which consisted of complete RPMI medium (Cat No. A1049101, Gibco, Waltham, Massachusetts, United States) supplemented with 50 ng/mL M-CSF (Cat No. 216-MC, R&D Systems, Minneapolis, Minnesota, United States) at a density of 1 × 10^6 cells/mL. The medium was replaced every 2-3 days, and the cells were allowed to differentiate into macrophages over a period of 7 days. We designated differentiated cells as WT-M cells.

### 2.7 Administration of WT-M through tail vein

The WT-M cells were injected into tumor-bearing mice by intravenous injection 14 days after tumor inoculation. After several days later, tumor tissues were harvested and analyzed that described in section 2.5.3 and 2.8-9.

### 2.8 Tumor dissociation

Tumors were isolated (described in section 2.5) and digested using Tumor Dissociation Kit (Cat. No. 130-096-730, Bergisch Gladbach, North Rhine-Westphalia, Germany). Tumors were cut into small pieces and incubated in an enzyme mix and placed in gentleMACS™ C Tube (Cat. No. 130-093-237, Bergisch Gladbach, North Rhine-Westphalia, Germany) attached upside down onto the gentleMACS™ Dissociator for 45 minutes. Cell suspension was then passed through a Falcon 70 µm cell strainer (Cat. No. 352350, Corning, New York, United States), washed with PBS twice and incubated in ACK Lysis Buffer (Cat. No.A1049201, ThermoFisher Scientific, Waltham, Massachusetts, United States) at room temperature for 10 minutes for removal of red blood cells.

### 2.9 Fluorescence Activated Cell Sorting (FACS)

Cells were washed with PBS and incubated in 5 uL of Human TruStain FcX™ (Fc Receptor Blocking Solution) (Cat. No. 422301, Biolegend, San Diego, California, United States), in 100uL FACS buffer (2% FBS in 1x PBS) for 10 minutes at room temperature, then incubate in Anti-CD45 Mouse Monoclonal Antibody (PE (Phycoerythrin)/Cy7®) (Cat. No. 304016, Biolegend, San Diego, California, United States) (1:500) for 30 minutes on ice. Samples are washed twice with FACS buffer prior to FACS.

## 3 Results

### 3.1 Preparation of oHA and characterization of CMCTS/oHA *in situ* hydrogels

As shown in Fig. 2a, by using sodium periodate, the cyclic structure of hyaluronic acid (HA) was oxidatively cleaved into an open chain and two aldehyde groups are produced from the vicinal diol units of the HA main chains, which eventually results in the oHA [23]. The formed dialdehyde group was confirmed by ^1^H NMR spectroscopy, where the peaks ranging from 4.9 to 5.1 ppm indicated the hemiacetal protons and vicinal hydroxyl of the dialdehyde group (Fig. 2b) [23, 24]. Consequently, newly generated dialdehyde group in oHA was able to be covalently cross-linked with amino groups in CMCTS via a Schiff base reaction [25, 26]. The formation of the CMCTS/oHA hydrogel was further verified by FT-IR (Fig. 2c). Compared to oHA, thestretching peak of dialdehyde group (-CHO) at 1730 cm^-1^ disappeared in the CMCTS/oHA hydrogel [27], while a characteristic absorption peak of dynamic imine bond emerged at 1590 cm^-1^ [28], which was demonstrated that CMCTS and oHA were clearly conjugated by Schiff-based reaction [29].

**Fig. 2.**
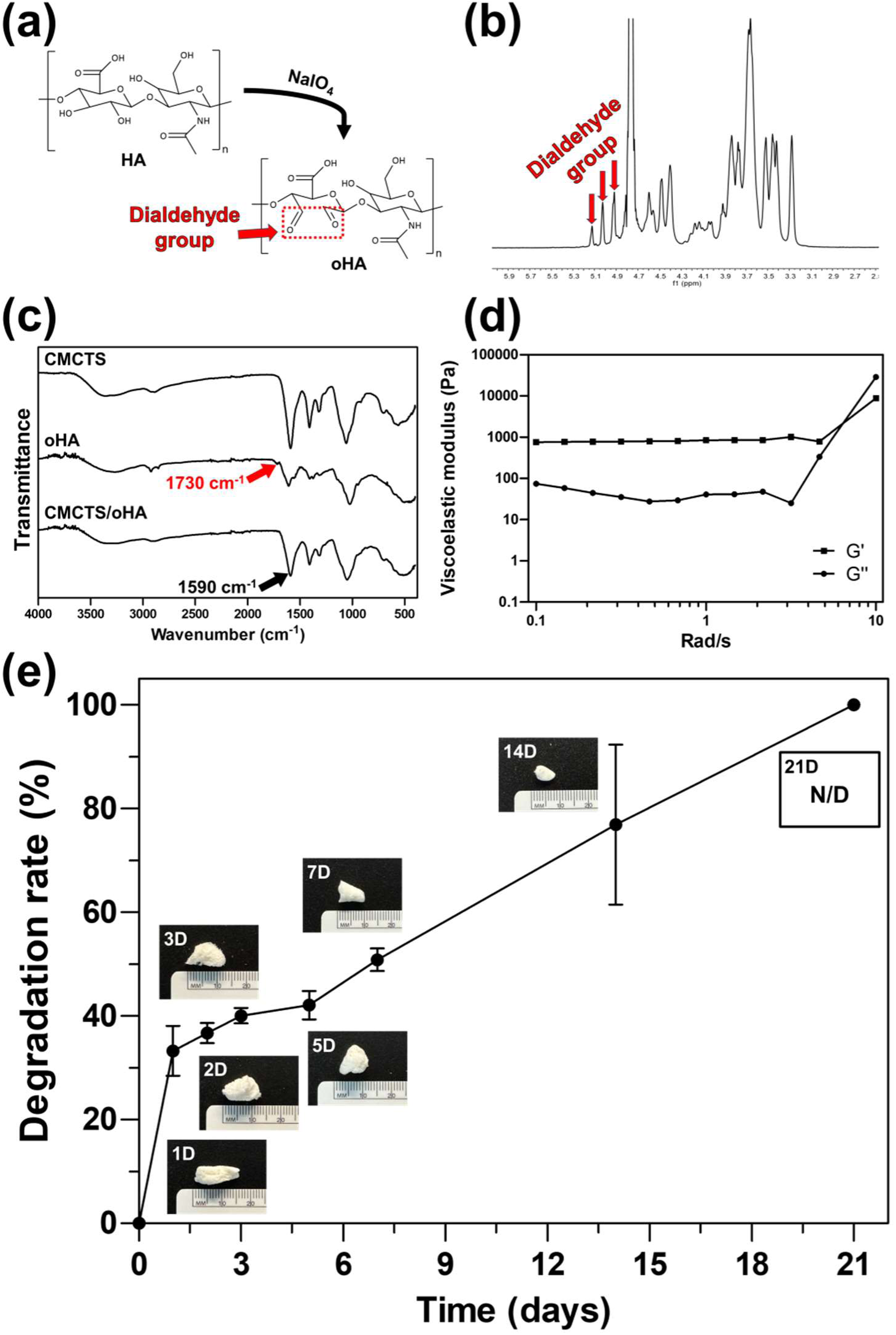
Characterization of CMCTS/oHA hydrogels. (a) Scheme of oHA synthesis by modification with NaIO4. (b) ^1^H NMR spectra of oHA indicated the presence of aldehyde groups. (c) FT-IR spectra of CMCTS, oHA composites and lyophilized CMCTS/oHA hydrogels. (d) Rheology analysis of storage modulus (G′), loss modulus (G″) of CMCTS/oHA hydrogels. (e) *In vitro* degradation of CMCTS/oHA hydrogels in PBS.

The transition of a hydrogel from sol to gel can be demonstrated by dynamic frequency scanning of the relative values between G’ and G’’. According to the dynamic frequency scanning curve, the values of G’ and G’’ remained comparatively steady and the energy storage modulus was higher than the loss modulus at the frequency below 5 Hz, suggesting the gel was dominated by elastic behavior in this range (Fig. 2d) [30]. However, when the frequency raised to 5 Hz, G’’ of hydrogels gradually became higher than G’, illustrating the system was closer to the solution state. CMCTS/oHA hydrogels as whole exhibited stable energy storage modulus and none of the G’’ values exceeded the G’ by two orders of magnitude, which indicated that they have a weak gelation behavior [26, 31].

The degradation behavior of CMCTS/oHA hydrogels was investigated by immersing them in PBS and recording their weight at specific intervals over time. As shown in Fig. 2e, the CMCTS/oHA hydrogel exhibited rapid degradation in the initial stage (within 24 hours), with a degradation rate of 30%. Subsequently, the degradation rate continued to increase at a relatively constant pace, leading to complete degradation of the hydrogel by 21 days of incubation. During the degradation process, significant decreases in sample volume were observed in images of freeze-dried hydrogels.

### 3.2 Preparation and characterization of high-density human-cancer cells loaded in CMCTS/oHA *in situ* hydrogels

First, the cell-free CMCTS/oHA hydrogel was directly extruded and exhibited a transparent, amorphous shape (Fig. 3a). Next, HepG2 encapsulated hydrogels were characterized. As shown in Fig. 3b and c, HepG2 cells were detected in the CMCTS/oHA hydrogel (red arrow), while the cell-free CMCTS/oHA revealed clear (yellow arrow). The high-density HepG2 cell-laden CMCTS/oHA was fabricated by tunnel mixing system (Fig. 3d) followed by extruded on well plate smoothly illustrated in Fig. 3e and f. On the day 0, HepG2 cells were encapsulated within the CMCTS/oHA hydrogel matrix (Fig. 3g). Over time, the cells began to condense starting from day 1, and the 3D tissue matrix became increasingly compact, resulting in highly condensed HepG2 tissue aggregates. It was also demonstrated rhodamine-labeled HepG2 cells demonstrated strong aggregate formation on 7 days of culture (Fig. S1).

**Fig. 3.**
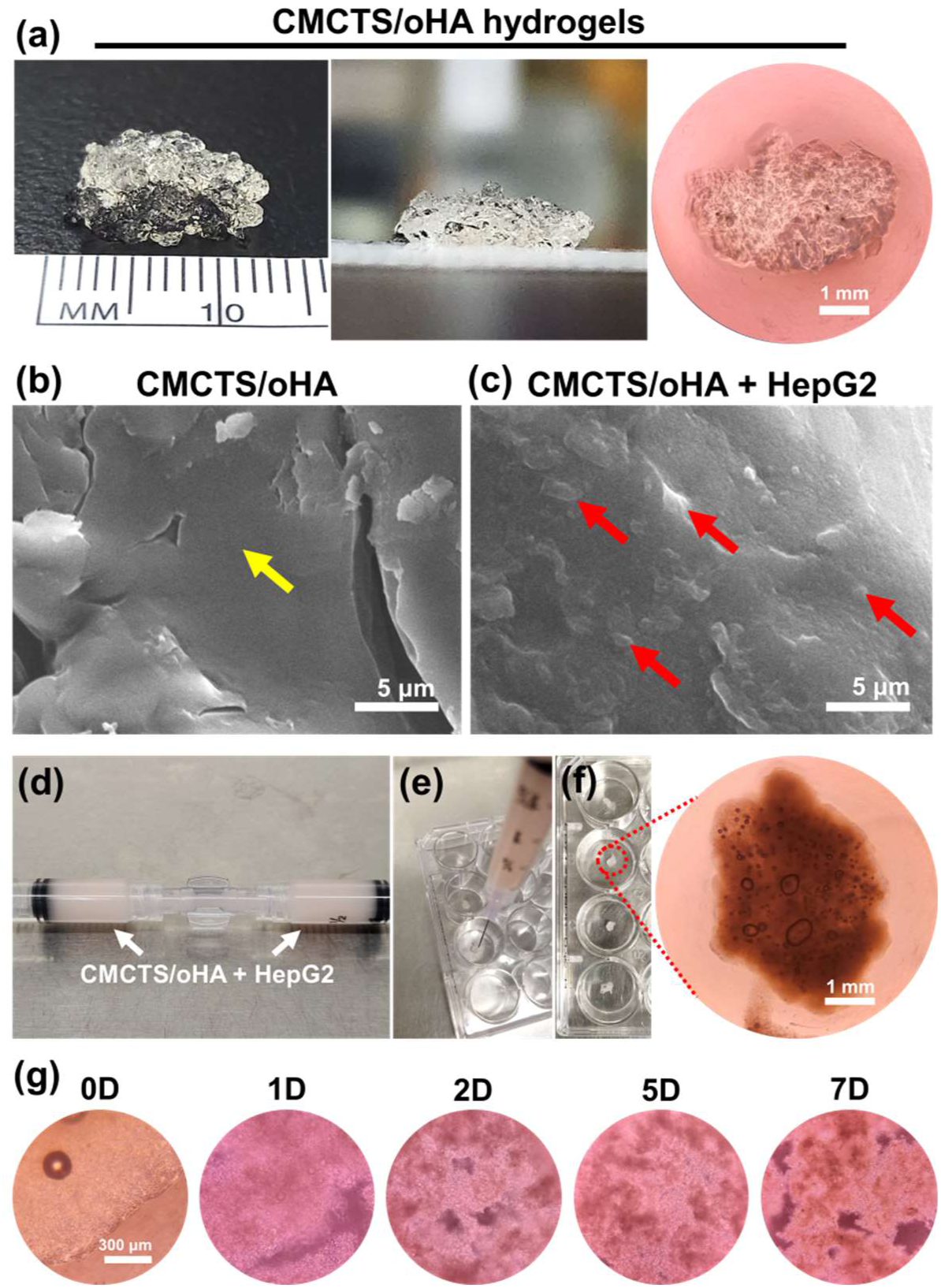
Injectability of CMCTS/oHA hydrogel and capacity to encapsulate high-density tumor cells. (a) Fabricated CMCTS/oHA hydrogel. Morphological analysis of the hydrogel before (b) and after (c) encapsulation of high-density HepG2 cells showed that cells (red arrows) were uniformly distributed in hydrogel (yellow arrows). (d) The tunnel mixing system enabled CMCTS/oHA gelation and thus direct introduction of high-density tumor cells without cross-linking agents, occurring at room temperature. The hydrogel encapsulating the tumor cells can be extruded directly through a needle using a syringe (e), and morphology of the hydrogel immediately after extrusion *in vitro* was observed using a microscope (f). (g) HepG2-containing CMCTS/oHA hydrogels were incubated *in vitro* for 7 days and gradually formed HepG2 tissue aggregates.

Also, HepG2 and Huh-7 cells were separately encapsulated in CMCTS/oHA hydrogel and cultured for 3 days (Fig. 4). The optical camera image indicated that the cell-laden CMCTS/oHA hydrogels, which initially paste status, but changed into a cell aggregate after 1 day of post-culture (Fig. 4a). Unfortunately, many cells were released down on plate during hydrogel degradation (Fig. 4b). On day 3 of culture, the 3D cell condensation showed high intensity of hoechst and rarely detectable dead cells (PI), indicating excellent cell viability (Fig. 4c). The morphology of cell matrix was confirmed by H&E staining and F-actin staining with nuclei. It indicated cells were connected closely as aggregate and condensed each other (Fig. 4d and e).

**Fig 4.**
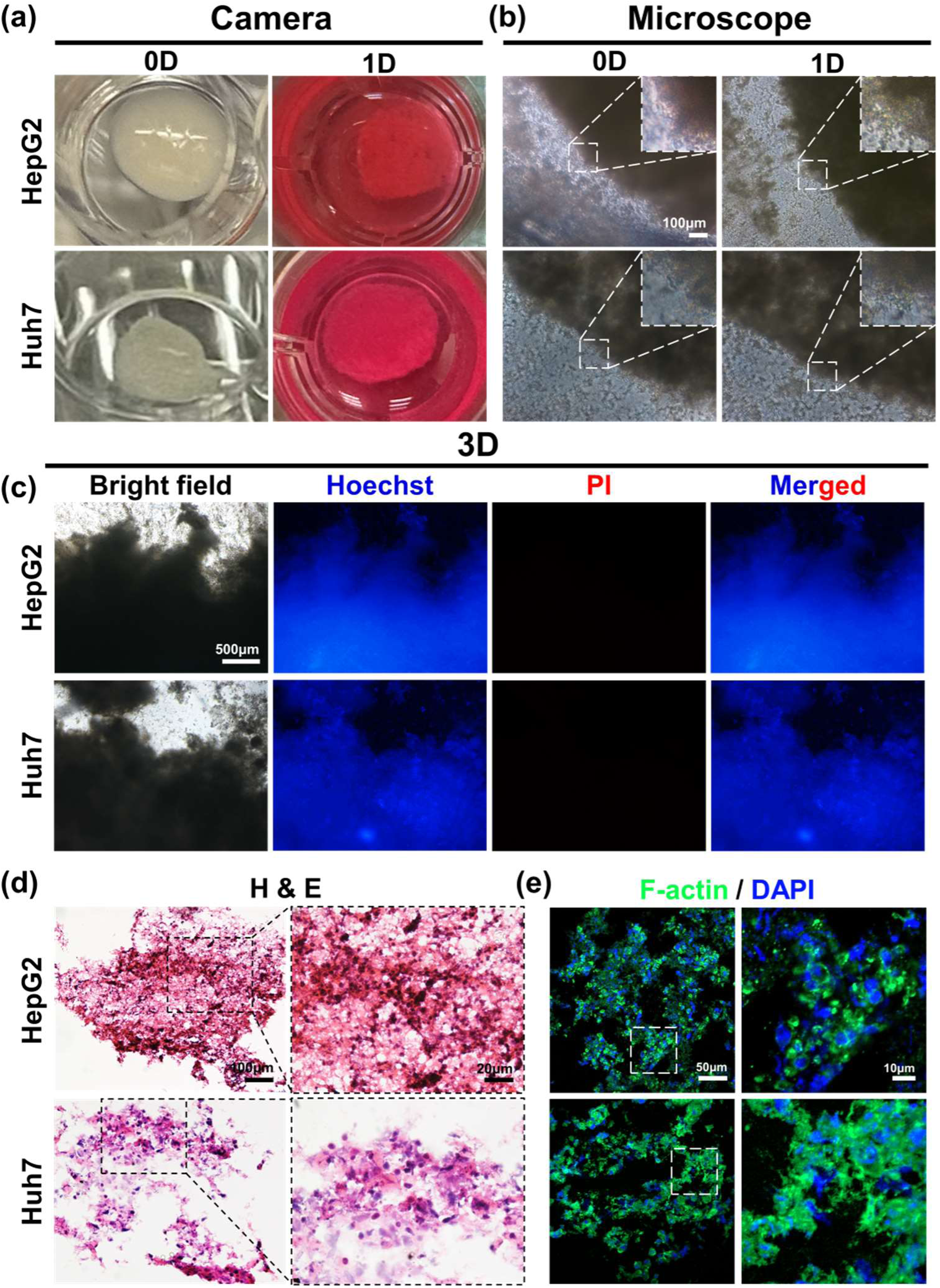
Cell viability of high-density HepG2 and Huh-7 cells encapsulated in CMCTS/oHA hydrogels. *In vitro* culture of high density HepG2 and Huh-7 cells encapsulated in CMCTS/oHA hydrogels for 3 days. Cells began to be released from the hydrogel from within the first day of *in vitro* culture (a-b). Cells are highly active, cohesive, and highly uniformly organized based on Hoechst 33342/PI assay (c), H&E staining (d) and F-actin staining (e) of cell-laden CMCTS/oHA hydrogels at 3 days of culture.

### 3.3 Degradation test of high-density human-cancer cells loaded in CMCTS/oHA *in situ* hydrogels via Balb/C mice subcutaneously

To assess the degradability of CMCTS/oHA hydrogels encapsulating HepG2 cells in the body, cell-laden hydrogels were directly injected subcutaneously into Balb/C mice. Transplants were obtained 1 week and 2 weeks post-injection (Fig. 5). Histological analysis revealed that HepG2 cells were embedded in CMCTS/oHA as single cells in 0D, and they did not form cell-to-cell connections (red arrow). 1 week post-injection, staining results showed well-preserved connective tissue in the native host group. Transplanted cells exhibited aggregation (blue arrow) but were closely connected to the native host tissue (green arrow). However, the cells did not form an extracellular matrix, which was attributed to the immune response from the Balb/C mice [32]. This was more clearly visualized 2 weeks post-injection, where the high-density HepG2 aggregates exhibited a strong connection with the host tissue in H&E and alcian blue staining.

**Fig. 5.**
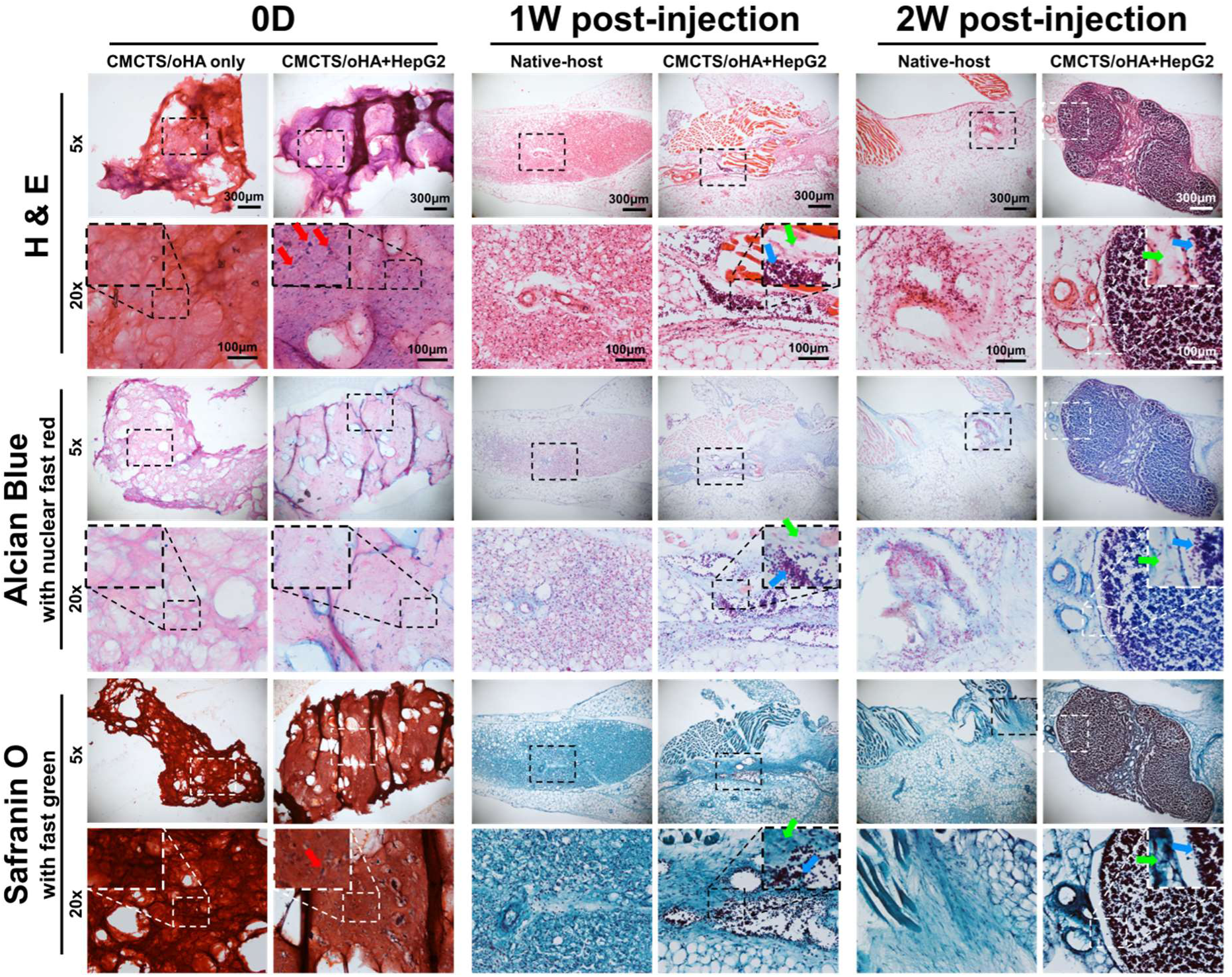
*In vivo* biocompatibility and biodegradability assessment of HepG2-encapsulated CMCTS/oHA hydrogels. High concentration HepG2-encapsulated CMCTS/oHA hydrogels were injected subcutaneously in Balb/C mice and their optical images were analyzed histologically 1 and 2 weeks after injection. Typical mouse subcutaneous tissues, such as adipose, muscle, and connective tissues, were visualized in all staining, with no remaining hydrogels. Tumor cells (blue arrows) were found present at all injection sites and were well integrated with the native tissue (green arrows).

Interestingly, safranin O with fast green staining indicated that the CMCTS/oHA hydrogel appeared strong red in the 0D group. However, the red color completely disappeared from the transplantation area 1 week post-injection. Result suggests that the hydrogel was spontaneously and completely degraded in the body, facilitating integration between the transplanted cells and host native tissue. Although no strong transplanted cell-to-cell connections and vasculature were found in the tumor area, high-density HepG2 cells established connections with the host native tissue (Fig. S2).

### 3.4 Generation of vascularized tumor model of high-density human-cancer cells loaded in CMCTS/oHA *in situ* hydrogels via SCID mice subcutaneously

To study neovascularization in tumor tissues, the CMCTS/oHA hydrogel loaded with high-density HepG2 cells was injected subcutaneously into SCID mice (Fig. S3) and histologically analyzed 2 weeks post-injection (Fig. 6). Histological staining of the HepG2-encapsulated *in situ* hydrogels before injection (0D) showed that the cells remained as single, free-floating entities within the hydrogel (Fig. 6a at 0D). 2 weeks after injection, solid tumors were observed at the injection site, exhibiting vascular interconnections (Fig. 6a at 2W). H&E and Alcian blue staining revealed that the HepG2 cells in the solid tumors formed large clusters, had a high nucleus-to-cytoplasm ratio, and displayed irregular nuclear shapes, confirming tumor formation. Safranin O staining indicated no residual red color in the tumor tissues, suggesting that the hydrogel was completely degraded. Notably, a “blood lake” morphology, characterized by clusters of red blood cells resembling vascular sprouts, was observed in all staining (asterisks in Fig. 6a). The presence of red blood cells was further confirmed by Wright-Giemsa staining (red asterisks in Fig. 6a). GPC3 and CD31 immunostaining also confirmed the presence of tumor capillaries, with CD31-positive staining indicating vascular structures in the tumor tissue (GPC3-positive staining) (Fig. 6b-d). The tumor tissues were surrounded by host adipose tissue, and the vasculature infiltrated the solid tumor. It was evident that the blood vessels originated from the host tissue in the mice, as no CD31 expression was detected in the surrounding tissue prior to transplantation (Fig. S4).

**Fig. 6.**
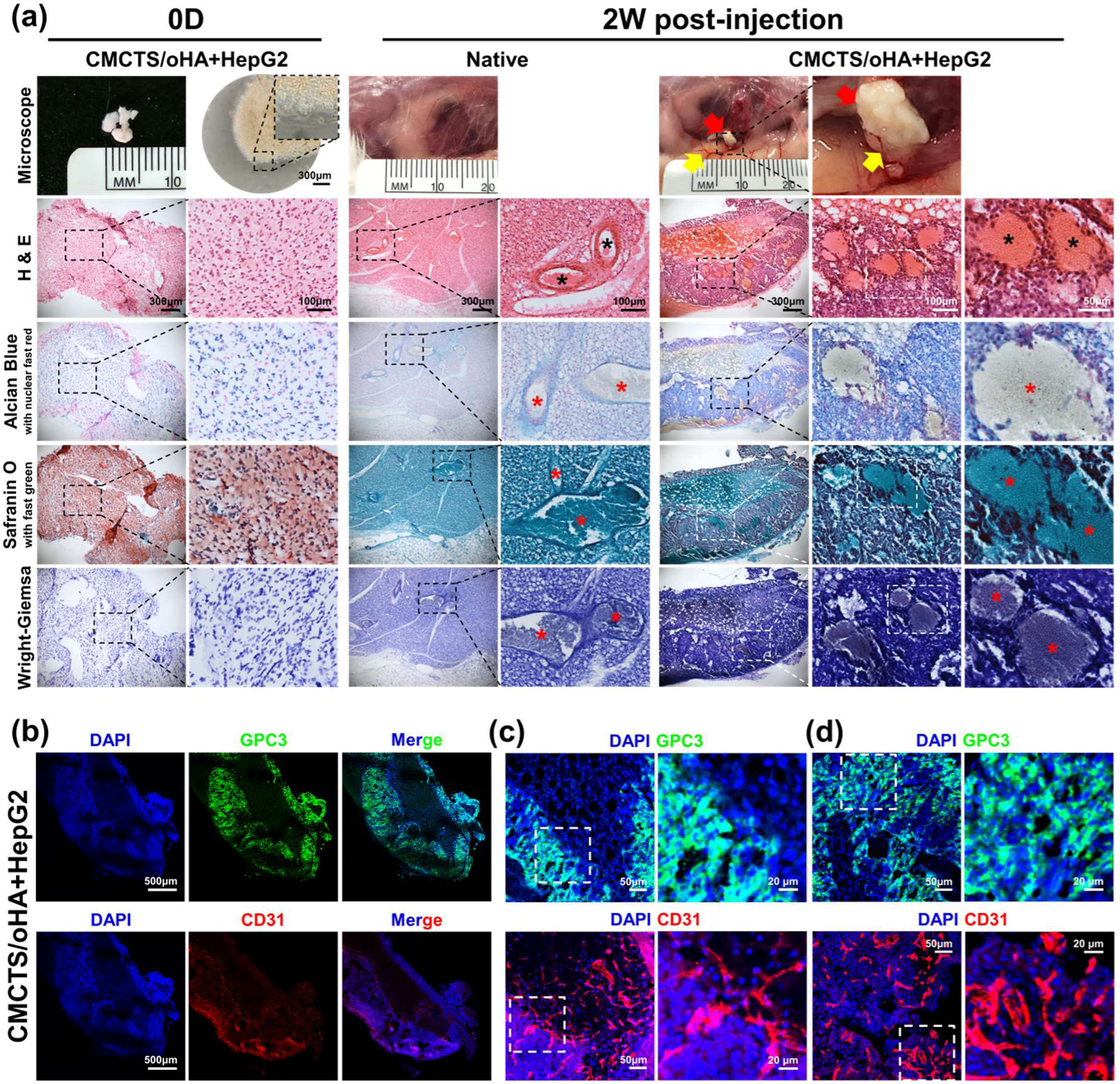
Vascularized tumor formation in immunodeficient SCID mice model. Highly concentrated HepG2-encapsulated CMCTS/oHA hydrogels were injected subcutaneously for 2 weeks (a). Mouse tissues such as adipose, muscle, and connective tissues, were visualized in all staining and no adverse tissue reactions such as necrosis or oedema were observed. Solid tumors (red arrows) were found and vessels (yellow arrows) were visualized on tumor surfaces. Histological staining of the tumor tissue showed the presence of red blood cell cluster (asterisks) internally within the tumor. IF staining images of GPC3 against HepG2 and CD31 against vasculature (b, c, and d). It clearly indicated that new vasculatures were infiltrated in solid tumor where closely connected with mouse-host adipose tissue.

To further investigate the vascularization of tumors after prolonged implantation, we examined the tumors 4 weeks post-injection of HepG2 and Huh-7 cell-encapsulated CMCTS/oHA hydrogels (Fig. 7). At the injection site, tumor tissues (red arrow) were observed to be tightly integrated with the mouse host tissue (white asterisk), and new vasculature was entangled with the tumor (yellow arrow) (Fig. 7a). H&E staining revealed robust vascular structures (yellow arrow), which included red blood cells (blue asterisk), indicating that blood was supplied within the tumor. CD31 immunofluorescence staining confirmed the presence of neovascularization by highlighting spindle-shaped tubular structures (Fig. 7b). A significant number of vessels were observed in various locations (Fig. 7b i-iv). Overall, our strategy demonstrated not only the straightforward formation of solid tumor tissue but also the generation of new vasculature within the tumor, enabling nutrient and oxygen circulation, which may enhance the effectiveness of cell therapy.

**Fig. 7.**
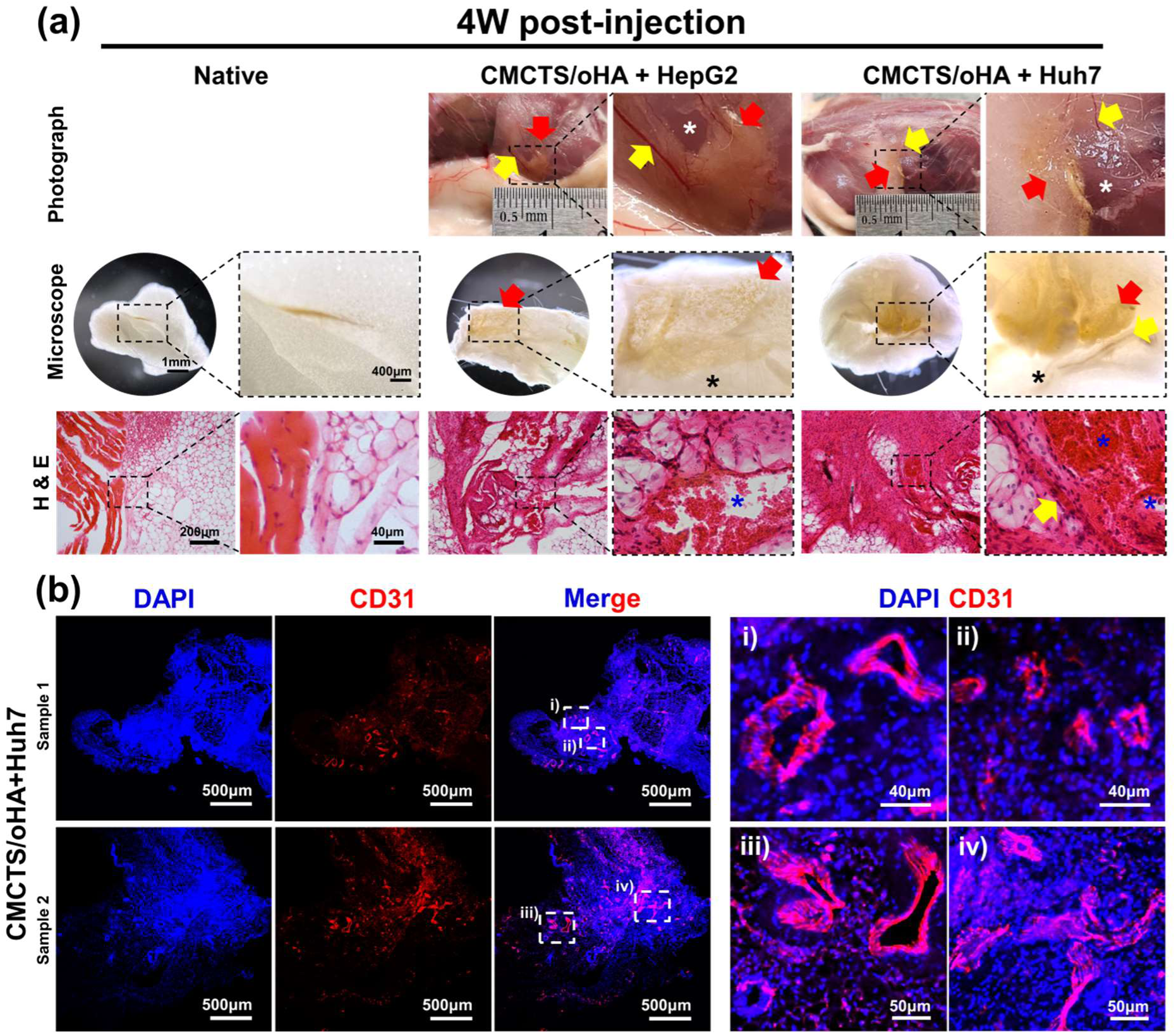
Vascularization of tumors in SCID mouse model for 4 weeks. High concentration of human liver cancer cells encapsulated CMCTS/oHA hydrogels were injected subcutaneously and analyzed (a). Tumors (red arrows) were generated on host muscle tissue (white asterisks) with vasculature (yellow arrows). The CD 31 IF staining showed that vascular lumen structures were strongly formed within solid tumors.

### 3.5 Adaptive immune cell therapy in the tumor model

To examine the infiltration of immune cells into vascularized tumors, we administered human immune cells through the bloodstream. Immune cell infiltration is a key criterion for effective cell therapy in clinical settings [33]. We hypothesized that the newly formed vasculature in solid tumors would facilitate this infiltration. Subcutaneous inoculation of HepG2 cells led to the formation of a visible tumor in the SCID mouse model (Fig. 8a), which grew significantly over the course of the experiment (Fig. 8b). The administration of human macrophages enhanced the accumulation of murine immune cells in the tumor, resulting in over a two-fold increase compared to the PBS injection group (Fig. 8c-d). This finding indicates a potential immune response against the tumor cells [34].

**Fig 8.**
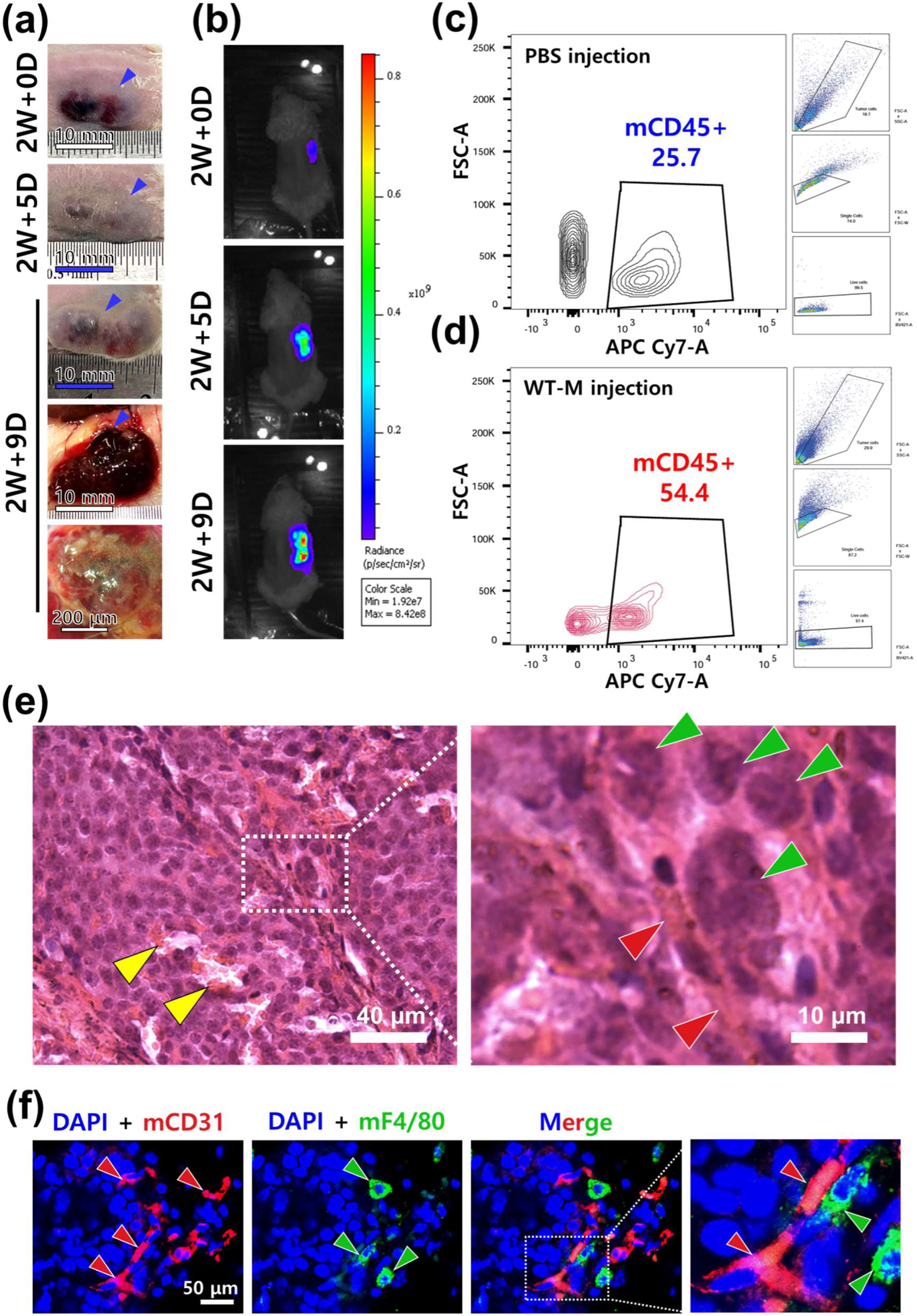
Adaptive cell therapy in vascularized tumors *in vivo*. Subcutaneous tumor formation in the SCID mouse model (a-b). Gross view of the tumor (a) and IVIS luminescence (b). 2 weeks post-injection of HepG2 cell-laden CMCTS/oHA hydrogels, human macrophages were intravenously administered via tail injection. “W” indicates weeks after tumor formation, and “D” indicates days after macrophage administration. Accumulation of host immune cells in the tumor due to adaptive cell therapy (c-d). Compared to PBS injection (c), macrophage administration (WT-M) resulted in a two-fold increase in mouse CD45+ immune cell accumulation in the tumor (d). H&E staining of the tumor tissue indicates perivascular macrophages (inlet) (e). The yellow arrow points to a sinusoidal vessel structure. The putative macrophage was adjacent to an arteriole structure. The red arrow indicated endothelial cells, and the green arrowhead indicated monocyte-like cells. Perivascular localization of murine macrophages was shown (f). Mouse CD31 (red) indicated endothelial cells, while mouse F4/80 (green) indicated murine macrophages neighboring the vasculature.

We confirmed the presence of putative human macrophages in the perivascular area of the tumor, which is consistent with previous findings that this region serves as an immunological hub within the tumor microenvironment [35]. The presence of murine macrophages in this area further supports the formation of this immune hub (Fig. 8f). Therefore, our findings demonstrate that a vascularized tumor model facilitates the infiltration of adaptive immune cell therapies and the establishment of perivascular immune hubs.

## 4 Discussion

Establishing xenograft human tumor models in mice is currently the most widely used approach for investigating antitumor therapies [36, 37]. These methods facilitate the rapid and extensive creation of *in vivo* tumor models. However, traditional injection of cell pellets does not guarantee consistent formation of solid tumors due to poor cell accumulation within the mouse [38]. Additionally, directly transplanted tumor cells often cannot survive without a supportive matrix [39]. Therefore, even when a uniform number of cells is transplanted, they may fail to adhere and proliferate at the target site, resulting in inconsistent tumor formation [40]. To address these issues, a stable carrier for cancer cells is essential for establishing *in vivo* tumor models that yield consistent therapeutic results. In our study, we employed the *in situ*-formed CMCTS/oHA hydrogel as a cancer cell delivery tool. This hydrogel possesses excellent biocompatibility and injectability, facilitating the formation of cell condensation (Fig. 3 and 4).

The CMCTS/oHA hydrogel effectively encapsulated and immobilized cells at the injection site during transplantation, fulfilling the critical need for reliable cancer cell carriers (Fig. S3). For generating a solid tumor model, it is essential that the hydrogel carrier is fully degradable and allows for cell release [41]. Additionally, the released cells must remain viable and be capable of spontaneously forming tumor tissue when the environment permits [42]. Our findings confirmed that the hydrogels were completely degraded post-injection, allowing for the generation of solid tumor clusters within the body (Fig. 5 and 6), which is consistent with the degradation profiles of the CMCTS/oHA hydrogel (Fig. 2d). The presence of cells within the hydrogel under cell culture conditions may enhance the dynamics of the hydrogel, thereby accelerating its degradation [43]. Consequently, our hydrogel demonstrated that cells were encapsulated within the hydrogel matrix and began to self-assemble upon release. Thus, our hydrogel system is effective for generating *in vivo* tumor models through spontaneous degradation *in situ*. However, it has not yet been demonstrated how tumor size and growth depend on the amount of injected cells, which requires further investigation.

Tumor cell survival and proliferation rely on the circulatory system for oxygen and nutrients, resulting in rapidly growing malignant tumors that are highly vascularized [44]. Our *in vivo* tumor model indicated that tumor cells and capillary endothelial cells were tightly integrated (Fig. 6 and 7). Tumors recruit blood vessels primarily through angiogenesis (the formation of new vessels from existing ones) and vasculogenesis (de novo vessel formation) [45]. These vascularization processes enable the development of extensive capillary networks; however, tumor vasculature often exhibits disordered structure and dynamics due to malignant growth and the overexpression of pro-angiogenic factors [46]. Additionally, some tumors, such as hepatocellular carcinoma, may utilize vascular mimicry, allowing tumor cells to form lumen-like structures independently of endothelial cell [47]. This phenomenon explains the presence of “blood lakes” in histological samples (Fig. 6). Ultimately, these vascularization pathways enable tumors to establish a dense network, facilitating an efficient blood supply within the dynamic tumor microenvironment (Fig. 7) [48].

One of the major obstacles in cell therapy is effectively delivering cells to tumors. The limited migration capability of T cell therapies, such as tumor-infiltrating lymphocytes (TILs), T cell receptor (TCR) T cells [49], and CAR-T cell therapy [50], hampers their application in solid tumors. As key components of the innate immune system with natural and superior tumor infiltration abilities, macrophage-based cell therapy appears to be a promising alternative for targeting solid tumors. When provided with appropriate cues, macrophages can be reprogrammed into a pro-inflammatory phenotype that engages in tumor killing [51]. *Ex vivo* polarized macrophages or the recent CAR-macrophages (CAR-M) therapy can be administered to patients for solid tumor treatment. Immune cell therapists claim that CAR-M cells have superior infiltration abilities compared to CAR-T cells [52], however, direct evidence of their migratory capacity remains limited [42, 43]. Our vascularized tumor model demonstrated that macrophages administered via injection could circulate and migrate within the tumors (Fig. 8). Their localization near blood vessels suggests the formation of cellular hubs, indicating potential crosstalk between immune cells and cancer cells [44]. While multiple clinical studies are underway for CAR-T cells, those for adoptive macrophage therapy remain limited. Establishing a reliable *in vivo* platform will facilitate more clinical trials for macrophage-based cell therapies.

Moreover, the spatial and temporal distribution of macrophages significantly impacts tumor prognosis [53]. Therefore, it is important to investigate the localization of adoptively transferred macrophages within the tumor microenvironment (TME), their migration, and their interactions with other cells, such as stromal, endothelial, and immune cells, to better understand how they exert their tumor-killing functions. Our fully vascularized tumor model provides a convenient platform to assess migration, spatial distribution, complex interactions with the TME, and potentially tumor-killing activity. Ultimately, combining vascularized tumor tissues with organs-on-chips and high-throughput imaging could advance chemical genetics screening for improved adoptive macrophage therapies. This approach addresses important parameters overlooked in previous macrophage therapy studies, such as persistence in tissues, clonal expansion of effector macrophages, and the invigoration of T cells through antigen presentation.

The limitation of the current study is the lack of molecular characterization of neovascularization in the tumor. A comparative analysis of our vascular structures with those of hepatocellular carcinoma patients would be important. Additionally, single-cell RNA sequencing may provide detailed phenotypes of endothelial cells, pericytes, and stromal cells. Nevertheless, we are confident that our evaluation will pave the way for the straightforward formulation of a fully vascularized tumor model in mice through simple injection of cells into the animal’s body.

## 5 Conclusions

In summary, our investigation successfully established an *in vivo* vascularized tumor mouse model using a homogeneous cross-linked *in situ* injectable CMCTS/oHA hydrogel that encapsulates high-density human liver cancer cells. The CMCTS/oHA hydrogels demonstrated excellent injectability and biocompatibility, with no significant adverse reactions observed after injection. The high-density human liver cancer cells encapsulated within the CMCTS/oHA hydrogels were completely degraded within 2 weeks post-implantation subcutaneously in mice. The released cancer cells aggregated at the injection site, leading to solid tumors that were tightly integrated with surrounding natural tissues. 4 weeks post-injection, mature vasculature was observed in these solid tumors. Additionally, injected macrophages, administered through the tail veins of the mice, were found near blood vessels within the tumors, indicating that the newly formed blood vessels were perfusable and could serve as a transport pathway for immune cells. Therefore, we believe that this *in vivo* vascularized tumor mouse model can provide valuable insights and feedback for strategies focused on macrophages, paving the way for future clinical applications in immune cell therapy against cancer.

## Author contributions

All authors have made contributions to this study work. Ziqi Huang, Yip Ming Tsun, and Chao Liang conducted the experiments, performed data analysis, and wrote original manuscript. Sangjin Lee and Zhenzhen Wu supported the experiments on material synthesis and structural analysis. Theo Aurich and Lu Liu assisted with *in vivo* experiments. Rio Ryohichi Sugimura and Sangjin Lee proposed valuable advice and provided financial support. All authors discussed the results and contributed to the manuscript preparation.

## Data availability

No datasets were generated or analysed during the current study.

