## Supplementary for "Homogeneously crosslinked *in situ* hydrogel enclosing high-density human-cancer cells promotes vascularized *in vivo* tumor modeling for immune cell therapy"

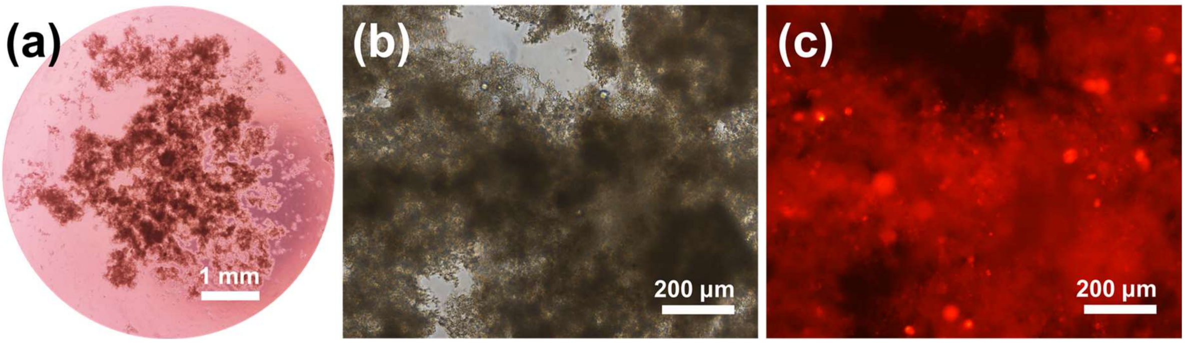


**Figure S1.** Microscope images of rhodamine-labeled HepG2 cultured in CMCTS/oHA hydrogels for 7 days *in vitro*. High-density HepG2 cells were formed aggregated (a) and exhibited strong cell-to-cell connection (b and c).

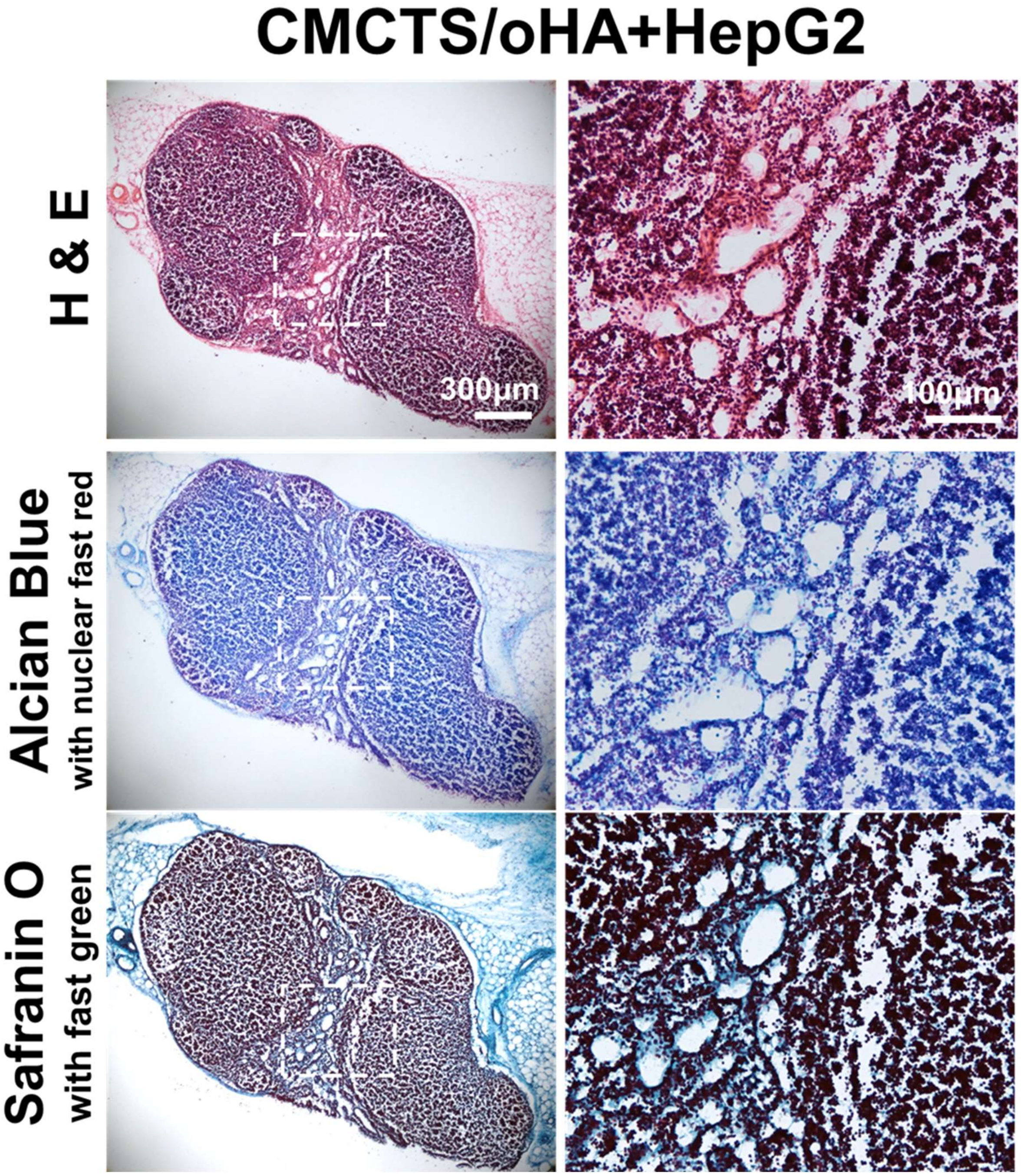


**Figure S2.** H&E, Alcian Blue and Safranin O staining of injection site after subcutaneous injection of high concentration HepG2-encapsulated CMCTS/oHA hydrogel over 2 weeks. Transplants were accumulated and connected with host-mouse tissue.

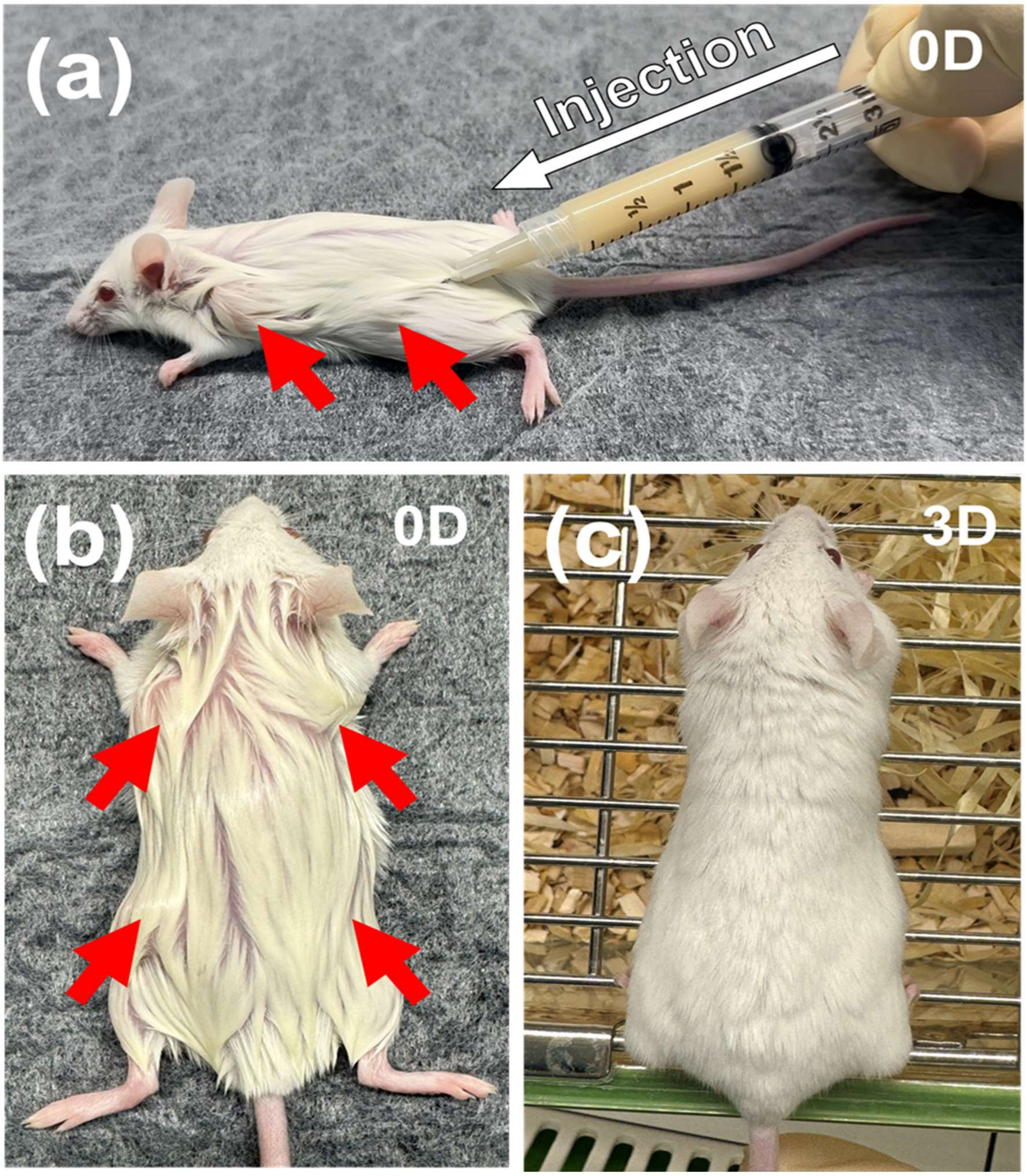


**Figure S3.** Administration procedure by subcutaneous injection. Hydrogel was injected subcutaneously on the back of mice without any incision (a). The injected hydrogel maintained the integrity of its volume in the immediate postoperative period (b), and the site volume receded after 3 days (c).

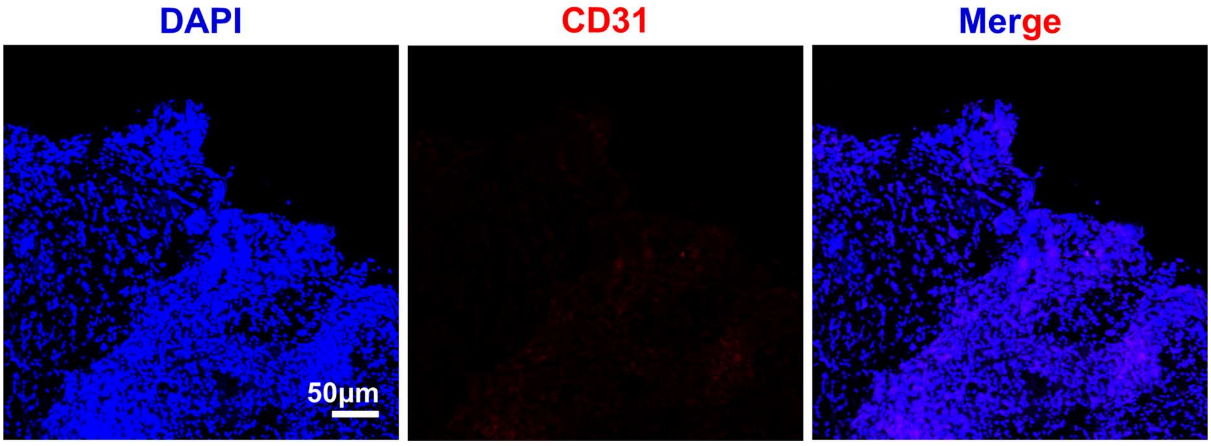


**Figure S4.** Immunofluorescence staining analysis of HepG2-encapsulated CMCTS/oHA hydrogel before implantation. CD31 showed no positive staining, demonstrating that the hydrogel was not pre-vascularized before injection.
